# FGF21 Response Varies by Sugar Type and is Associated with Body Weight, Dietary Added Sugar, and Neural Signaling in Humans

**DOI:** 10.1101/2021.05.29.446318

**Authors:** Alexandra G. Yunker, Jasmin M. Alves, Shan Luo, Brendan Angelo, Alexis DeFendis, Trevor A. Pickering, Kay Jann, John R. Monterosso, Kathleen A. Page

**Affiliations:** Division of Endocrinology, Department of Medicine, Keck School of Medicine, University of Southern California, Los Angeles, CA 90089; Diabetes and Obesity Research Institute, Keck School of Medicine, University of Southern California, Los Angeles CA 90089; Department of Psychology, University of Southern California, Los Angeles, CA 90089, USA; Department of Preventive Medicine, Keck School of Medicine, University of Southern California, Los Angeles, CA 90089; Mark & Mary Stevens Neuroimaging & Informatics Institute, Keck School of Medicine, University of Southern California, Los Angeles, CA 90089

**Keywords:** Fibroblast growth factor 21 (FGF21), sucrose, fructose, glucose, obesity, dietary added sugar, brain, humans

## Abstract

Fibroblast growth factor 21 (FGF21) is a liver-derived hormone that regulates energy homeostasis. In humans, few studies have investigated whether FGF21 may act to suppress sugar intake and influence eating behavior, and the effects of adiposity on post-ingestive FGF21 regulation of appetite are unknown. Here, we demonstrate among two cohorts of healthy, young adults that acute oral fructose and sucrose compared to glucose lead to greater circulating FGF21. Moreover, high compared to low dietary added sugar intake is associated with greater sucrose-stimulated FGF21 among participants with healthy weight but attenuated in people with overweight and obesity. In addition, our study is the first to demonstrate associations between circulating FGF21 and neural signaling following an acute sucrose load among humans with healthy weight. Collectively, our results suggest that these potential compensatory relationships between sucrose-stimulated circulating FGF21, habitual sugar intake, and post-ingestive brain responses may be altered among adults with overweight and obesity.

**Significance Statement:** Animal models have established FGF21 as an autoregulator of sweet consumption, but few studies have examined post-ingestive FGF21 effects in humans. In this report, we demonstrate a compensatory relationship between sucrose-stimulated FGF21 and high dietary added sugar intake through a potential liver-to-brain negative-feedback cycle among healthy, young adults. Notably, our findings also suggest that humans with overweight and obesity may have altered FGF21 neuroendocrine signaling.

## Introduction

Obesity rates have risen dramatically over the last three decades, posing a significant challenge to public health (1). A growing body of evidence has linked sugar consumption to increased rates of weight gain, risk for obesity, and development of chronic disease (2–4). The common dietary sugars, glucose, fructose, and sucrose (a disaccharide composed of equal parts glucose and fructose, commonly known as “table sugar”), are absorbed and metabolized differently by the body, and prior studies by our group and others have shown that these sugars have differential effects on hormones involved in glucose homeostasis, suggesting that each have distinct physiological consequences that may subsequently impact endocrine regulation of appetite, feeding behavior, and body weight (5–8).

Fibroblast growth factor 21 (FGF21) is a liver-derived hormone that regulates energy homeostasis and plays a critical role in several metabolic and physiological processes in both rodents and humans (9, 10). In particular, FGF21 is an important regulator of carbohydrate appetite and sweet taste preference (11). Notably, in humans, FGF21 has been shown to robustly increase after acute oral fructose (12), but its rise is attenuated in response to glucose (12–14), likely due to differences in hepatic metabolism between the two monosaccharides. Interestingly, fructose-stimulated FGF21 levels were shown to be increased in adults with metabolic syndrome compared to healthy participants (12), however, the effects of BMI (i.e., healthy weight, overweight, and obesity) on the dynamic FGF21 response to different types of sugars among healthy, young adults is currently unknown.

In addition to the previous findings showing fructose-linked increases in FGF-21 secretion, Søberg and colleagues recently demonstrated that plasma FGF21 also increased markedly after an oral sucrose load in healthy weight adults, and that sweet-disliking participants have elevated fasting FGF21 levels (15). Moreover, studies indicate that FGF21 gene variates are associated with a preferential increase in dietary carbohydrate intake (16, 17), particularly added sugar and candy consumption (15), and that FGF21 selectively reduces appetite for simple sugars without changing intake of other macronutrients or overall calorie intake (11). Mechanistic studies in rodents and non-human primates demonstrate that FGF21 reduces sweet taste preference through its effects on striatal dopamine signaling (18) and hypothalamic neurons (11, 19). Collectively, these findings are suggestive of FGF21 acting as a down-regulator of subsequent sweet consumption through its effects on the central nervous system and suggest a role of FGF21 in nutrient-specific appetite regulation. While previous studies suggest that FGF21 may negatively autoregulate sugar consumption through a liver-to-brain feedback cycle (11, 15), no study has examined how circulating FGF21 affects neural responsivity following sucrose ingestion in humans. Moreover, prior studies suggest a large variability of the FGF21 response between subjects (12, 20), indicating the importance of identifying how biological (i.e., body weight) and environmental (i.e., dietary sugar intake) factors might affect the FGF21 response to acute sugar ingestion.

To address these gaps in knowledge, in this report we examined three questions: (1) How is BMI status associated with the dynamic FGF21 response to oral fructose vs glucose and sucrose vs glucose among two cohorts of human subjects? (2) Does the FGF21 response to oral sucrose vary by habitual dietary sugar intake (i.e., high vs low dietary sugar) among young adults with healthy weight, overweight or obesity? (3) Do circulating FGF21 levels affect neural responsivity in response to oral sucrose, and does BMI influence this relationship?

## Results and Discussion

Thirty-eight participants (19 male, 19 female) aged 16-25 years with a BMI range between 19.1-45.4 kg/m^2^ completed the Brain Response to Sugar I pilot study (Experiment 1). Sixty-nine adults (29 male, 40 female) aged 18-35 years with a BMI range between 19.2-40.3 kg/m^2^ completed the larger Brain Response to Sugar II study (NCT02945475) (Experiment 2). Participant characteristics for each cohort are provided in **Table 1**. In each of these studies, participants arrived after a 12h overnight fast and we assessed plasma FGF21 before and after participants consumed either a standard 75g glucose (Experiment 1 and 2), 75g fructose (Experiment 1), or 75g sucrose (Experiment 2) load, mixed with 0.45g of non-sweetened zero calorie cherry flavoring for palatability, dissolved in 300mL of water. In Experiment 1, blood samples were collected at baseline (0min) and approximately 75min post-drink. In Experiment 2, blood samples were collected in accordance with a standard oral glucose tolerance test (OGTT): at baseline (0min), 10min, 35min, and 120min post-drink (21, 22). Trajectories for post-prandial plasma FGF21 at each time point in Experiments 1 and 2 are shown in **Figure S1**. In addition, in Experiment 2 we also measured dietary added sugar intake through multiple 24h dietary recalls assessments and brain response to oral sucrose using perfusion magnetic resonance imaging (MRI). For detailed study overviews, see **Materials & Methods**.

**Table 1.** Participant characteristics for Experiment 1 and Experiment 2.

| Experiment 1 (N=38) |  |  |
| --- | --- | --- |
| Variable | Mean (SD) or N (%) | Range |
| BMI (kg/m<sup>2</sup>) | 28.5 (7.1) | 19.1-45.4 |
| Age (years) | 21.6 (2.0) | 16.1-25.2 |
| Sex | Males: 19 (50%)<br>Females: 19 (50%) |  |
| BMI Status | Healthy Weight: 20 (53%)<br>Obesity: 18 (47%) |  |
| Fructose drink condition |  |  |
| Baseline FGF21 (pg/ml) | 255.8 (269.6) | 10.0-1231.0 |
| Stimulated FGF21 (pg/ml) (75min) | 327.9 (278.0) | 10.0-1240.4 |
| Glucose drink condition |  |  |
| Baseline FGF21 (pg/ml) | 283.1 (329.9) | 10.0-1641.0 |
| Stimulated FGF21(pg/ml) (75min) | 256.9 (304.5) | 10.0-1561.5 |
| Experiment 2 (N=69) |  |  |
| Variable | Mean (SD) or N (%) | Range |
| BMI (kg/m<sup>2</sup>) | 27.0 (7.1) | 19.2-40.3 |
| Age (years) | 23.2 (3.7) | 18.2-34.5 |
| Sex | Males: 29 (42%)<br>Females: 40 (58%) |  |
| <sup>a</sup>BMI Status | Healthy Weight: 25 (36%)<br>Overweight: 23 (33%)<br>Obesity: 21 (30%) |  |
| Sucrose drink condition |  |  |
| Baseline FGF21(pg/ml) | 284.5 (247.8) | 5.5-1192.8 |
| Stimulated FGF21 (pg/ml) (120min) | 611.1 (478.9) | 3.2-2644.8 |
| Glucose drink condition |  |  |
| Baseline FGF21(pg/ml) | 292.3 (270.6) | 6.8-1132.7 |
| Stimulated FGF21 (pg/ml) (120min) | 286.3 (245.7) | 7.4-1249.5 |

<sup>a</sup>Percentages may not be equal due to rounding to nearest whole number.

### FGF21 response is greater following fructose and sucrose vs glucose ingestion

In Experiment 1, we examined the FGF21 response to oral fructose vs glucose at ∼75 min relative to baseline since these blood sampling points were shared by all participants. While baseline levels of plasma FGF21 were not different between the fructose and glucose conditions [t(38)=0.71, p=0.48], we found that change in circulating plasma FGF21 levels from baseline to ∼75min were greater in response to acute fructose compared to glucose ingestion in the whole cohort, adjusted for age, sex, and BMI status [F(1,37)=8.52, p=0.01] (**Table S1, Figure 1**). These findings are complementary to those presented by Dushay and colleagues, who found that serum FGF21 levels rose robustly following fructose ingestion, while FGF21 did not markedly increase following an oral glucose load (12). In contrast to fructose, only a small fraction of an oral glucose load is metabolized by the liver, and these differential circulating FGF21 responses are likely explained by differences in hepatic fructose and glucose metabolism (12). Furthermore, in a recent review, von Holstein-Rathlou et al. noted that FGF21 may have evolved as a protective endocrine mechanism in order to negatively autoregulate sugar intake within healthy physiological limits, and the potential toxic effect of elevated levels of fructose intake on the liver may help to explain the higher FGF21 response to fructose relative to glucose consumption (23). However, it is important to note that the study design of Experiment 1 did not allow us to capture peak post-prandial FGF21 response after fructose or glucose ingestion, which has been consistently reported to occur at 120min in humans (12, 14, 23, 24). Søberg et al. first verified that sucrose consumption in humans leads to an increased production of FGF21 (15), and in Experiment 2 we captured their previously reported peak timepoint (120min) for FGF21 response to sucrose ingestion in humans (15). We found that while baseline levels of FGF21 were not different between the sucrose and glucose conditions [t(64)=0.02, p=0.99], the change in circulating FGF21 levels from baseline to 120min was greater in response to acute sucrose compared to glucose ingestion in the whole cohort, adjusted for age, sex, and BMI status [F(1,64)=266.54, p<0.0001] (**Table S2, Figure 1**). Notably, a prior study in rodents demonstrated that sucrose, compared to fructose and glucose, ingestion produces the highest levels of plasma FGF21 (11). However, we were not able to statistically compare rises in FGF21 between Experiment 1 and 2 due to differences in timing of blood sample measurements. Future studies that examine dynamic FGF21 response to different types of sugars should investigate whether these previous findings in rodents translate to humans by directly comparing circulating FGF21 after an oral sucrose vs fructose load.

**Figure 1.**
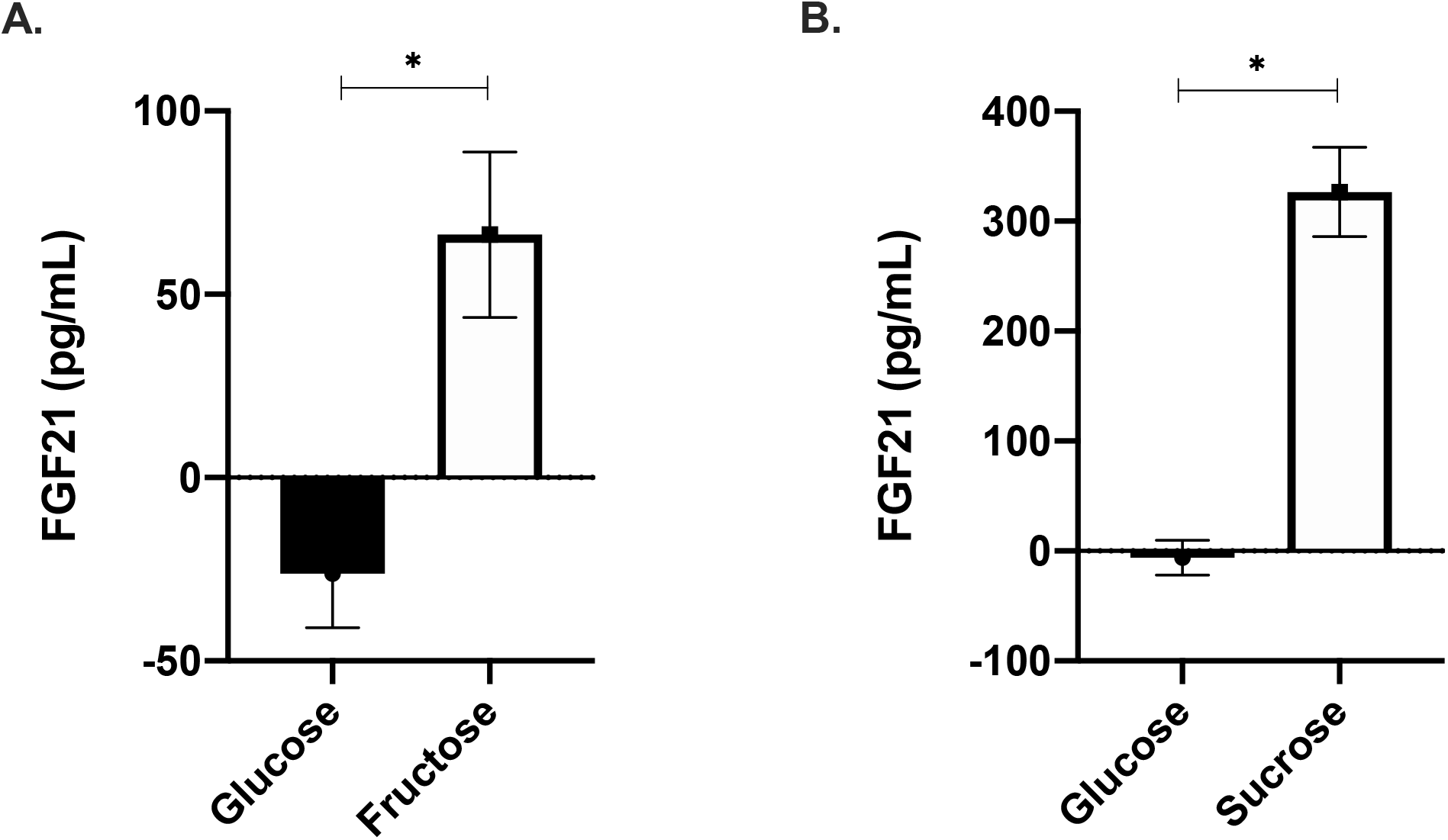
Stimulated plasma FGF21. Following acute A) glucose and fructose (Experiment 1, N=38); and B) glucose and sucrose consumption (Experiment 2, N=69), among the two cohorts. Stimulated-FGF21 was calculated as values at 75min minus baseline in Experiment 1, and as values at 120min minus baseline in Experiment 2. Data are expressed as raw/unadjusted mean ± SEM for visual purposes, but all statistical analyses were based on cubic root transformed FGF21 values and adjusted for covariates. See also Figure S1 and Tables S1 and S2.

In models stratified by BMI status in Experiment 1, participants with healthy weight had a significant increase in FGF21 levels from baseline to ∼75min after fructose vs glucose ingestion [F(1,19)=5.58, p=0.03], while individuals with obesity had a non-significant trend towards greater circulating FGF21 from baseline to ∼75min after fructose vs glucose ingestion [F(1,17)=2.97, p=0.10] (**Table S1**). Given the small sample size in our pilot study and the limited time of blood sampling measurements, additional studies are necessary to confirm these findings. In Experiment 2, FGF21 response from baseline to 120min was higher after sucrose relative to glucose consumption in all BMI groups: in participants with healthy weight [F(1,22)=106.35, p<0.01], overweight [F(1,22)=83.17, p<0.01], and obesity [F(1,18)=79.02, p<0.01], adjusted for age and sex (**Table S2**). This is the first study to show that acute oral sucrose, relative to glucose, stimulates a robust rise in circulating FGF21 among humans with a range of BMI. A previous report demonstrated that healthy weight adults experience increases in circulating FGF21 in response to an acute sucrose load over 300min, with peak FGF21 response occurring at 120min (15), and our current findings indicate that individuals with overweight and obesity also have potent sucrose-evoked increases in plasma FGF21 after 120min post-ingestion. Given that the participants with overweight and obesity in our cohort did not have a diagnosis of diabetes or other metabolic complications, future investigators should examine if overweight/obesity, in the presence of endocrine comorbidities (i.e., metabolic syndrome, diabetes), leads to altered sucrose-stimulated FGF21 secretion.

### High compared to low dietary added sugar intake is associated with greater FGF21 response to acute sucrose ingestion among individuals with healthy weight

In Experiment 2, we administered repeated 24-hour dietary recalls (an average of five dietary recalls over approximately 2 months per participant) to determine how habitual dietary added sugar consumption was related to the FGF21 response to acute sucrose ingestion. For the purpose of this study, we analyzed the amount of added sugar in the diet as a percent of total calories, and we classified participants as either high or low dietary added sugar consumers based on the World Health Organization’s dietary recommendations: ≥10% added sugar was considered a high amount of added sugar and <10% was considered a low amount of added sugar (25). In our cohort, percent calories from dietary added sugar intake ranged from 1.6-29.3%, with 42 low-added-sugar (<10%) and 27 high-added-sugar (≥ 10%) consumers. There were no differences in habitual dietary added sugar intake (i.e., low or high) between individuals with healthy weight, overweight, or obesity [F(2,73)=0.47, p=0.63]. Dietary added sugar use was not associated with peak sucrose-stimulated FGF21 levels among the cohort as a whole (β=0.76, 95% CI: -0.25-1.77, p=0.14); however, among individuals with healthy weight, high vs low dietary added sugar intake was associated with a greater peak FGF21 response to oral sucrose, adjusted for covariates (β=1.90, 95% CI: 0.47-3.34, p=0.02) (**Table S3, Figure 2**). While similar patterns were observed in overweight and obese groups, the difference in peak FGF21 response to oral sucrose in high vs low dietary added sugar consumers was attenuated and non-significant (**Table S3, Figure 2)**. Overall, these data indicate that high (vs. low) dietary sugar consumption is associated with a robust FGF21 response to an acute sucrose ingestion among individuals with healthy weight, and this response is attenuated in individuals with overweight/obesity. These data are in line with the concept that the liver responds to chronic high sugar diets in a compensatory fashion by markedly increasing FGF21 in response to an acute sugar load, and FGF21 in turn acts as negative endocrine regulator of sugar consumption (11, 15, 18, 23). While our correlative findings need to be followed up with experimental studies to determine causal mechanisms, the notion that high dietary sugar intake increases FGF21 production as a protective endocrine mechanism is supported by data in mice showing that a high sucrose diet increased hepatic FGF21 production and FGF21 sensitivity in brown adipose tissue, which attenuated weight gain (26). Correspondingly, human studies showed that when compared to a control diet, short-term high carbohydrate feeding led to a striking 8-fold increase in plasma FGF21 levels, which affected glucose and lipid homeostasis in healthy male volunteers (27). Based on our preliminary findings, it is possible that individuals with overweight and obesity might have diminished sucrose-evoked FGF21 compensatory signaling for dietary added sugar use, which could in turn contribute to a vicious cycle of high habitual added sugar intake and obesity, but this possibility requires additional investigation.

**Figure 2.**
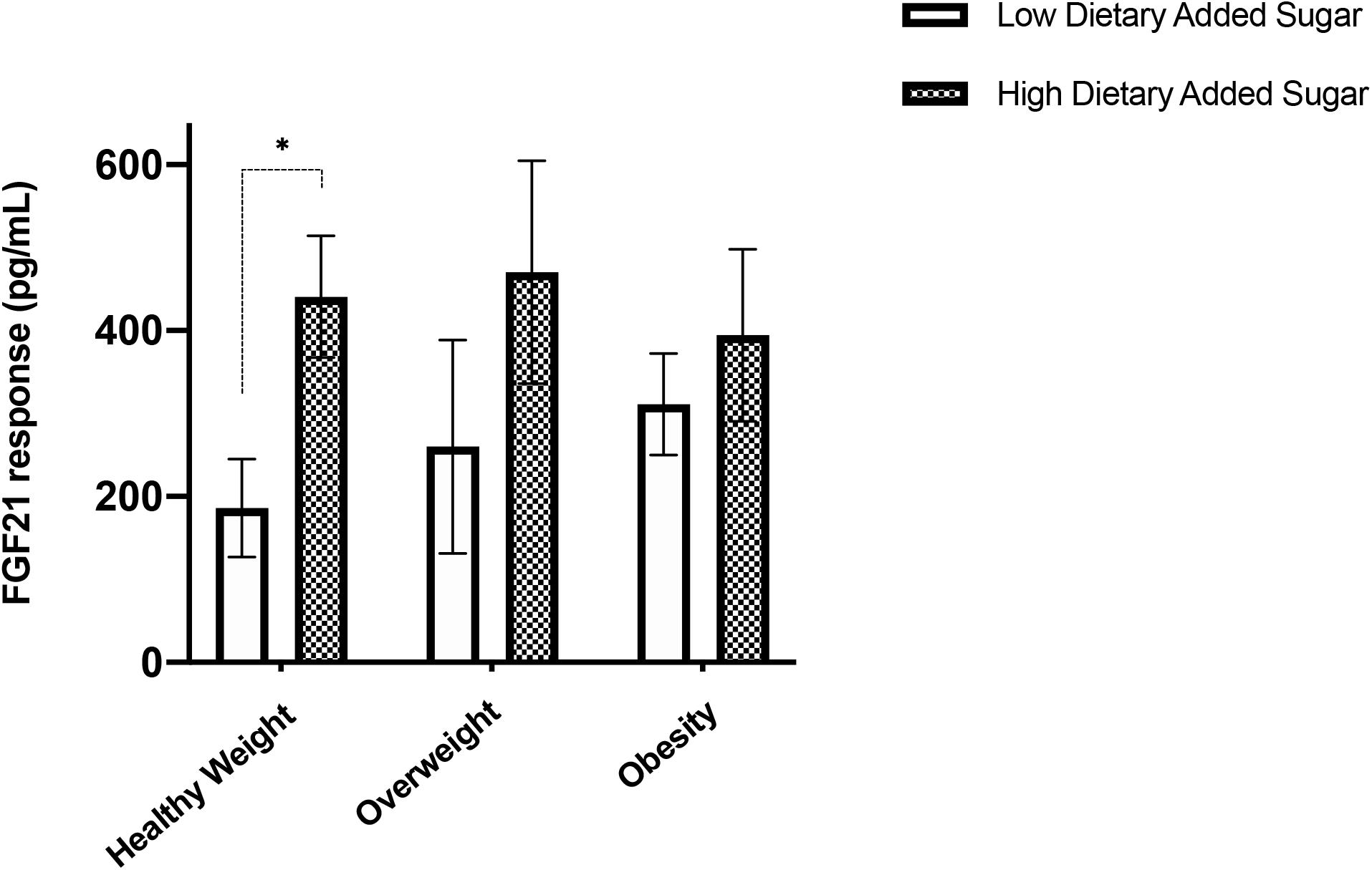
Relationship between high vs low dietary added sugar intake and sucrose-evoked FGF21 response. High compared to low dietary added sugar intake was associated with a greater FGF21 response to acute sucrose ingestion among people with healthy weight, but results were attenuated in participants with overweight and obesity. Data are expressed as raw/unadjusted mean ± SEM for visual purposes, but all statistical analyses were based on cubic root transformed FGF21 values and adjusted for covariates. See also Table S3. *Indicates significant difference in FGF21 response between high vs low dietary added sugar consumption at p<0.05.

### Negative associations between FGF21 and neural response to sucrose ingestion is driven by individuals with healthy weight

*A priori* brain regions-of-interest (ROI) included areas implicated in sweet taste processing and/or were selected based on evidence demonstrating FGF21 neural expression: dorsal striatum, hippocampus, hypothalamus, insula, and nucleus accumbens (11, 18, 19, 28–32). We calculated total area under the curve (AUC)_35_ for plasma FGF21 and cerebral blood flow (CBF) based on Arterial Spin Labeling (ASL) perfusion MRI in each ROI using the trapezoid method in order to quantify the FGF21 and neural responses to sucrose ingestion across the first phase 35min time period when both neuroimaging and blood sampling were simultaneously performed. Prior studies have demonstrated that, among healthy adults, a post-prandial state leads to a decreased CBF response in the brain areas involved in food intake regulation, including the hypothalamus, insula, hippocampus, and striatum (6, 33–37), which may be indicative of neural satiety signaling. Furthermore, altered post-prandial CBF signaling in these neural regions is associated with obesity (33, 34, 38–42). In the current study, we found a negative association between FGF21 AUC_35_ and dorsal striatum CBF AUC_35_ in response to sucrose ingestion (β=-7.66, 95% CI: -14.48 to -0.85, p=0.03), adjusted for covariates (age, sex, BMI status, global mean CBF (mCBF), insulin AUC_35_, and glucose AUC_35_). Similar trends were observed in the hippocampus, insula, and nucleus accumbens (**Table S4**). In models stratified by BMI status and adjusted for covariates, we observed a negative association between FGF21 AUC_35_ and dorsal striatum CBF AUC_35_ in response to sucrose ingestion among individuals with healthy weight (β=-15.70, 95% CI: -25.98 to -5.41, p=0.01), but not with overweight (β=-4.00, 95% CI: -19.72-11.72, p=0.62) or obesity (β=-12.45, 95% CI: -27.42-2.51, p=0.13) (**Figure 3**). Correspondingly, we also found that individuals with healthy weight had negative associations between FGF21 AUC_35_ and hippocampal CBF AUC_35_ in response to sucrose (β=-20.60, 95% CI: -36.79 to -4.42, p=0.02), but these associations were attenuated among individuals with overweight (β=-17.87, 95% CI: -36.93-1.19, p=0.08), and the associations were nonsignificant, but notably in the opposite direction, in individuals with obesity (β=1.34, 95% CI: -14.37-17.05, p=0.87) (**Figure 3, Table S5**). To the best of our knowledge, this is the first report to investigate associations between circulating FGF21 and neural responses to sucrose ingestion in humans. Our current findings support the hypothesis raised by previous investigators that circulating FGF21 may regulate sugar consumption through a liver-to-brain negative-feedback loop (11, 15). Here we provide evidence in line with their prediction, showing a negative association between circulating FGF21 and neural responses to sucrose ingestion in the dorsal striatum and hippocampus among healthy young adults. The dorsal striatum plays a critical role in motivational desire for food (43), and the hippocampus acts to integrate post-nutritive signals in order to suppress appetite (44–46). Prior studies have shown that excess weight gain and overweight/obesity are associated with impairments in striatal and hippocampal signaling (47–52). In the current study, when we stratified the results by BMI group, we found that participants with healthy weight, but not overweight or obesity, had a negative association between circulating FGF21 levels and striatal and hippocampal responses to acute sucrose ingestion. These data indicate that individuals with overweight and obesity might have altered FGF21-linked neuroendocrine appetitive signaling, which could lead to subsequent increases in appetite and eating behavior. Future longer-term studies in humans are needed to elucidate the role of FGF21 in neuroendocrine regulation of appetite and body weight among individuals with varying levels of adiposity.

**Figure 3.**
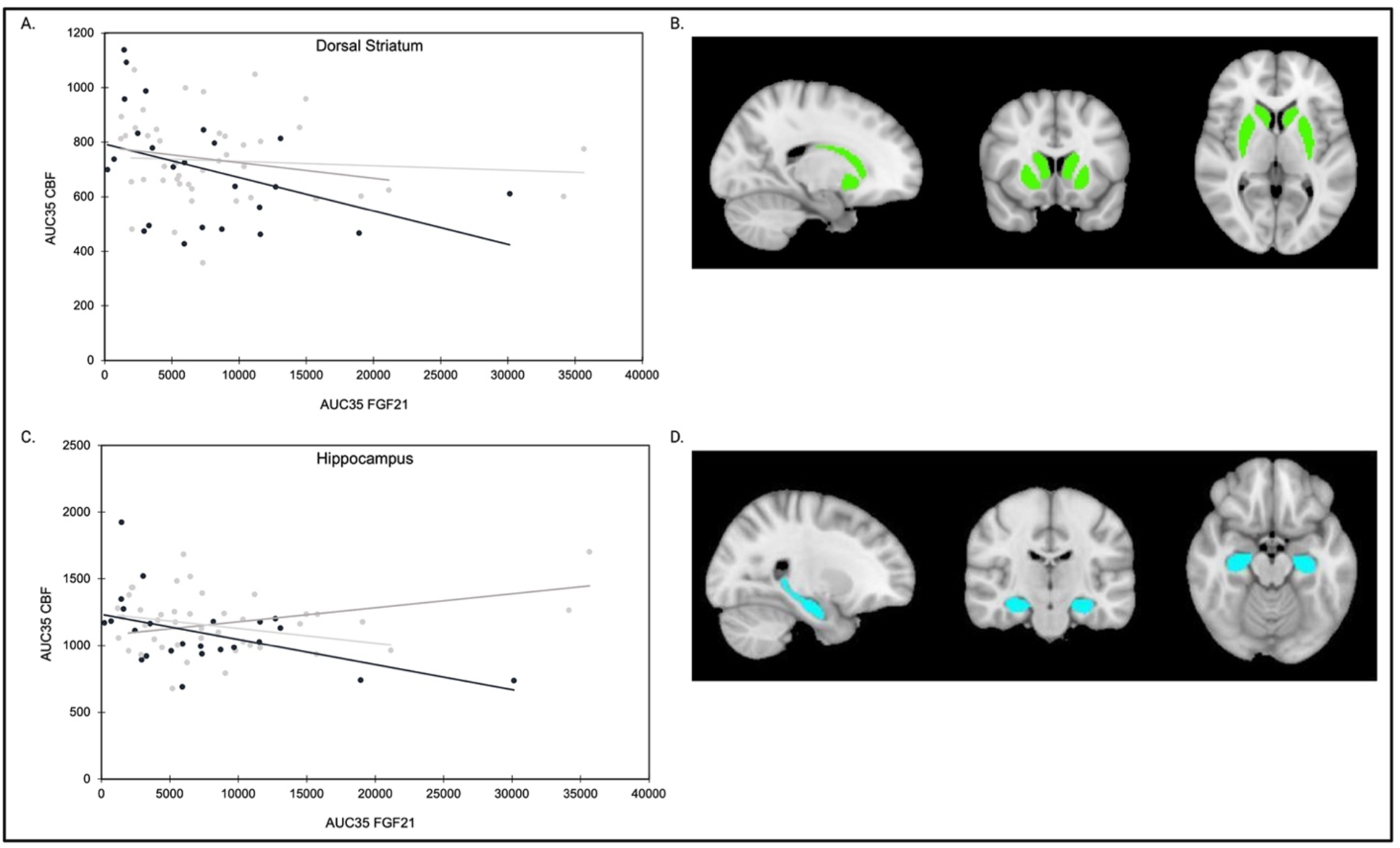
Associations between sucrose-stimulated circulating FGF21 and brain signaling. Correlational graphs show relative change in plasma FGF21 AUC_35_ from pre- to post-sucrose load was significantly negatively associated with A) dorsal striatal and C) hippocampal CBF AUC_35_ response to sucrose ingestion among young adults with healthy weight (black), but not with overweight (light gray) or obesity (dark gray). Data in panels A) and C) are expressed as raw/unadjusted AUC_35_ values for visual purposes, but all statistical analyses were based on cubic root transformed FGF21 values and adjusted for covariates. Region of interest (ROI) mask of B) dorsal striatum [Total voxels = 2392; center voxel Left = -26, 2, 2; center voxel Right = 28 2, 2] and D) hippocampus [Total voxels = 1033; center voxel Left = -28, -20, -16; center voxel Right = 30, -22, -14.] derived from the Harvard-Oxford subcortical atlas. See also Tables S4 and S5.

It is important to note that this was a preliminary study and the first to investigate associations between circulating FGF21 and neural signaling in humans. Our findings set the stage for future research that should examine relationships between dynamic FGF21 and brain responses across longer testing periods and among additional human populations (i.e., children/adolescents and older adults) to examine liver-to-brain signaling developmentally and across the lifespan. Future investigation into the role of FGF21 neural signaling on sweet taste preference, food cravings, and eating behavior is also warranted. Furthermore, given that previous reports examining the neural FGF21 receptor complex have largely been centered on targeting networks in the hypothalamus and nucleus accumbens (18, 19, 23, 30, 53–55), our findings have important implications for future in vitro and rodent studies that should consider the dorsal striatum and hippocampus as potential neural regions for FGF21 expression and signaling.

In summary, our study among young adult humans reveals three important findings: (1) circulating FGF21 response is greater following acute fructose and sucrose compared to glucose ingestion; (2) high vs low dietary added sugar intake is associated with greater FGF21 responses to an oral sucrose load among individuals with healthy weight, but not overweight or obesity; and (3) circulating FGF21 is negatively associated with striatal responses to oral sucrose (see **Figure 4**, conceptual model). These findings provide important insights into the potential compensatory relationship between high added sugar intake and sucrose-stimulated FGF21 signaling. Our study is the first to report a potential liver-to-brain negative-feedback cycle between circulating FGF21 and dorsal striatal and hippocampal responses to acute sucrose ingestion among adults with healthy weight, and our results suggest that FGF21 neuroendocrine regulation of appetite may be altered among individuals with overweight and obesity. This work supports the emerging role of FGF21 as an endocrine regulator of sugar intake in humans and the importance of considering how individual factors, including adiposity and/or diet, may influence physiological mechanisms underlying nutrient-specific appetite. Overall, this work implicates targeted dietary interventions as a promising strategy to benefit gut-brain feedback mechanisms, eating behavior, and metabolic health.

**Figure 4.**
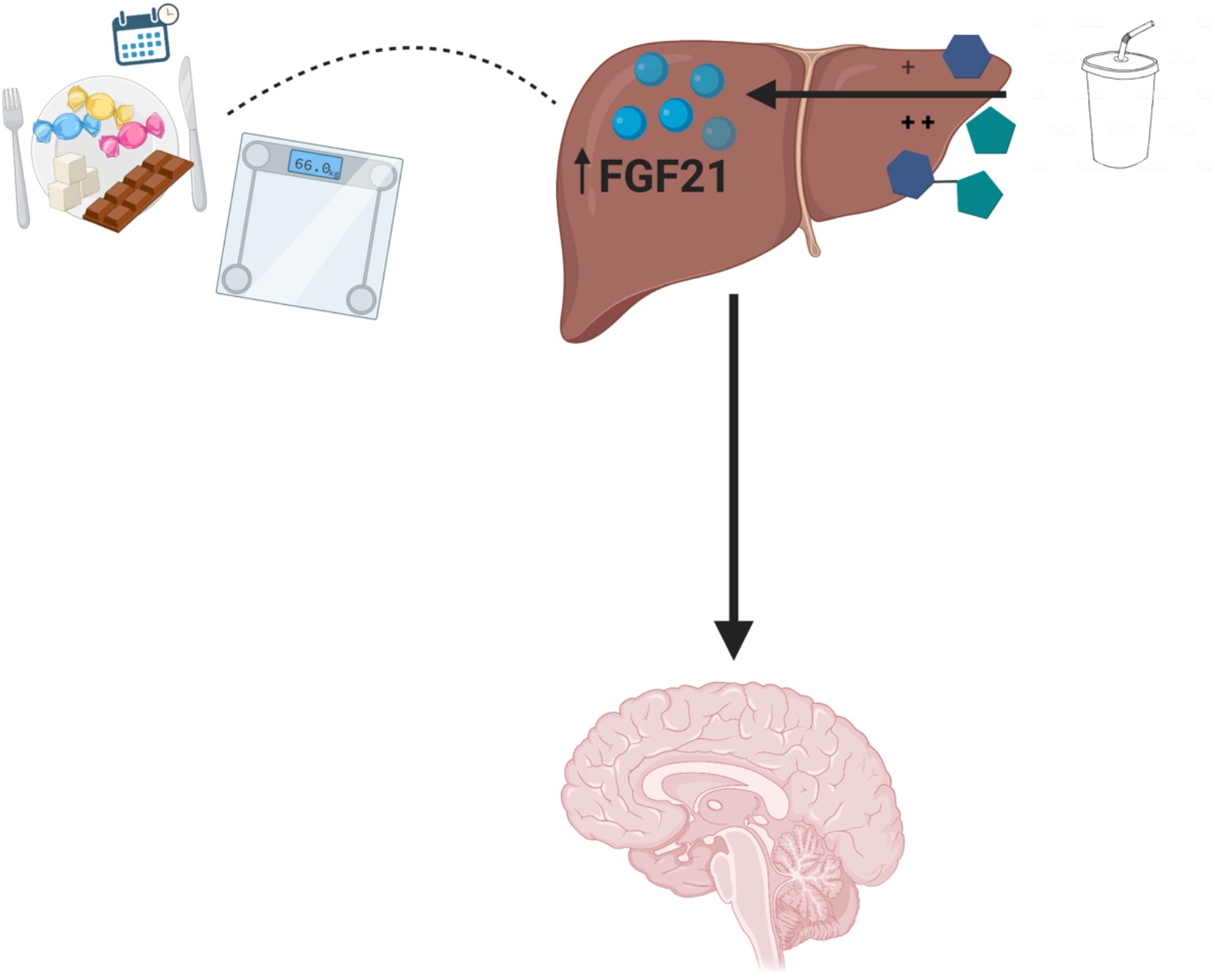
Conceptual Model. FGF21 response is greater following acute fructose and sucrose vs glucose. Circulating FGF21 is associated with post-ingestive brain responses, and FGF21 signaling may be impacted by both habitual sugar intake and obesity.

## Materials & Methods

### Experiment 1 Study Overview

The data reported here is part of the Brain Response to Sugar I pilot study. Participants gave written informed consent to all experimental procedures approved by the Institutional Review Board of the University of Southern California (IRB #HS-09-00395). Recruitment occurred between January 2013 and November 2015. Participants were right-handed, nonsmokers, non-dieters, not on any medication (except oral contraceptives), and with no history of diabetes, fructose intolerance, eating disorders, illicit drug use, or other medical diagnoses. Participants were asked to maintain their typical diets and physical activity levels throughout the study.

All the visits were conducted in the morning after a 12h overnight fast. Participants underwent a screening visit during which height was measured to the nearest 0.1cm using a stadiometer and weight to the nearest 0.1kg using a bioelectrical impedance analysis scale (Model no. SC-331S, TANITA Corporation of American, Inc.), and BMI was calculated as weight in kilograms divided by height in meters squared. Participants were split into two BMI status groups (healthy weight: 18.5-24.9 kg/m^2^ or obesity: ≥ 30 kg/m^2^) based on Centers for Disease Control and Prevention (CDC) criteria (56). In addition, a 75g fructose tolerance challenge was also performed; only individuals who reported no significant gastrointestinal discomfort (i.e., bloating, nausea, flatulence, diarrhea) on a questionnaire administered 1h after the ingestion of the fructose load were included in the study in order to limit confounding effects of fructose malabsorption. Based on this criteria, ten participants were excluded from participating in the study due to fructose intolerance.

Participants arrived at the University of Southern California at approximately 8:00AM for their study visits. The time interval between the two sessions was between 2 and 30 days. The order of the drinks was randomized using a computer-generated sequence. Participants and experimenters were blind to the drink provided during the study sessions. Participants consumed either a standard 75g glucose or 75g fructose load, mixed with 0.45g of non-sweetened zero calorie cherry flavoring (Kraft Foods Kool-Aid® Unsweetened Cherry Drink Mix) for palatability, dissolved in 300mL of water. The drinks were equicaloric (300 kcal each). During the study visits, blood samples were collected at baseline (0min) and at approximately 75min post-drink to measure plasma FGF21. Female subjects underwent study visits during the follicular phase of their menstrual cycles.

### Experiment 2 Study Overview

The data reported here is part of the larger Brain Response to Sugar II study examining neuroendocrine responses to high-reward foods (NCT02945475). This sub-study consisted of an initial screening visit, an oral glucose tolerance test (OGTT), and an oral sucrose tolerance test (OSTT). Participants provided written informed consent compliant with the University of Southern California Institutional Review Board (IRB #HS-09-00395). Recruitment occurred between July 2016 and March 2020. The Brain Response to Sugar II within-participant randomized crossover trial included four drink conditions (see online digital repository for full trial protocol (57)), but the data analyzed for this manuscript included two drink conditions (i.e., sucrose and glucose) to test the a priori hypothesis that oral sucrose and glucose would have differential effects on circulating FGF21 responses. For the Brain Response to Sugar II study Consolidated Standards of Reporting Trials (CONSORT) flow diagram see online digital repository (57).

All the visits were conducted in the morning after a 12h overnight fast. During the screening visit, we assessed eligibility for participation in the study and demographic information and anthropometric measurements were obtained. Participants were right-handed, nonsmokers, weight-stable for at least 3 months prior to the study visits, non-dieters, not on any medication (except oral contraceptives), and with no history of diabetes, eating disorders, illicit drug use, or other medical diagnoses. Height was measured to the nearest 0.1cm using a stadiometer and weight to the nearest 0.1kg using a bioelectrical impedance analysis scale (Model no. SC-331S, TANITA Corporation of American, Inc.), and BMI was calculated as weight in kilograms divided by height in meters squared. Participants were split into three BMI status groups (healthy weight: 18.5-24.9 kg/m^2^, overweight: 25-29.9 kg/m^2^, and obesity: ≥ 30 kg/m^2^) based on Centers for Disease Control and Prevention (CDC) criteria (56).

Participants arrived at the Dornsife Social and Behavioral Phlebotomy Lab at University of Southern California at 8:00AM for their study visits. The time interval between the two sessions was between 2 and 30 days. The order of the drinks was randomized using a computer-generated sequence. Participants and experimenters were blind to the drink provided during the study sessions. Participants consumed either a standard 75g glucose or 75g sucrose load, mixed with 0.45g of non-sweetened zero calorie cherry flavoring (Kraft Foods Kool-Aid® Unsweetened Cherry Drink Mix) for palatability, dissolved in 300mL of water. The drinks were equicaloric (300 kcal each). During the study visits, blood samples were collected in accordance with a standard OGTT: at baseline (0min), 10min, 35min, and 120min post-drink to measure plasma FGF21, glucose, and insulin. Female subjects underwent study visits during the follicular phase of their menstrual cycles.

As part of the larger Brain Response to Sugar II study, participants also underwent Magnetic Resonance Imaging (MRI) sequences in the Dornsife Cognitive Neuroimaging Center of University of Southern California during each study visit starting at approximately 9:00AM. Baseline MRI sequences included a T1 structural scan (for anatomical registration) and an arterial spin labelling (ASL) scan, followed by participants exiting the scanner to consume the test drink within two min (in order to reduce variability in timing of drink effects). After consuming the drink, participants re-entered the scanner and underwent two additional ASL sequences at approximately 5 and 30min post-drink ingestion. A food cue task measuring blood oxygen level dependent (BOLD) signaling was also collected as a part of the larger study, but not included in the current analysis.

### Metabolite and Hormone Analysis

Plasma FGF21 (pg/ml) was measured using a human FGF21 ELISA kit (Millipore, Billerica, MA). In Experiment 2, plasma glucose (mg/dl) was measured enzymatically using glucose oxidase (YSI 2300 STAT PLUS Enzymatic Electrode-YSI analyzer, Yellow Springs Instruments) and plasma insulin (pg/ml) was measured via Luminex multiplex technology (Millipore, Billerica, MA).

### Habitual Dietary Added Sugar Intake Assessment

In Experiment 2, diet was assessed using the multipass 24-hour dietary recall, which is a validated method that provides detailed information on food and beverages consumed over the previous 24-hour period (58, 59). Each dietary interview was administered by a trained staff member, wherein volunteers were asked to recall all food and drinks items (including meals and snacks) that they ingested during the previous 24-hours. In order to account for potential daily variations in dietary intake, 24-hour recalls were captured on both weekdays and weekend days. After the dietary recalls were obtained, the data was analyzed using the Nutritional Data System for Research (NDSR) software v.2018, developed by the Nutrition Coordinating Center, University of Minnesota, Minneapolis, MN, USA (60). Dietary recalls were also assessed for plausibility and quality using the method described by Jones et al. (49), and using this method, 359 dietary recalls were included in the analysis (an average of 5 dietary recalls, over the course of approximately 2 months, per participant) and 5 recalls were excluded. The output from the software provided intake of overall calories and the breakdown of macronutrients.

### MRI Imaging Parameters and Analysis

In Experiment 2, neuroimaging data were collected using a 3T Siemens MAGNETOM Prismafit MRI System, with a 32-channel head coil. After initial localizers, a baseline pulsed arterial spin labeling (pASL) scan was acquired followed by a high-resolution 3D magnetization prepared rapid gradient echo (MPRAGE) sequence (TR=1950ms; TE=2.26ms; bandwidth=200Hz/pixel; flip angle=9°; slice thickness=1mm; FOV=224mm×256mm; matrix=224×256), which was used to acquire structural images for multi-subject registration. The ASL sequences used the QUIPSS-II method (61). A proximal inversion with a control for off-resonance effects (PICORE) mode was employed to provide high labeling efficiency (62). The ASL sequences were acquired along with one M0 image with following parameters: field of view (FOV) = 192 mm, matrix = 64 × 64, bandwidth = 2232 Hz/Pixel, slice thickness = 5 mm, interslice spacing = 0 mm, Repetition time (TR) = 4000 ms, echo time (TE) = 30 ms, flip angle = 90°, TI1/TIs/TI2 = 700/1,800/1,800 ms, label duration = 1675 ms, slab thickness = 100mm, in-plane resolution = 3 × 3 mm^2^. The timing of the inversion pulses (TI) were optimized to reduce intravascular signal intensity at 3 T (63). The duration of the ASLs acquisition was 8 min and 18 seconds.

Pulsed ASL data was collected using magnetically tagged arterial blood. Motion was corrected (64) and slice timing correction used 53ms for CBF travel time. Tagged and untagged images were subtracted to obtain perfusion-weighted images. Bayesian Inference for ASL (BASIL) toolbox (https://fsl.fmrib.ox.ac.uk/fsl/fslwiki/BASIL), was used to analyze the perfusion data. Mean CBF across the whole brain and specifically in the brain regions-of-interest (ROI) were extracted. All ROI were bilateral and anatomically defined using the Harvard-Oxford cortical and subcortical structural atlas found in FSL, except the hypothalamus which is not included in the atlas and was defined bilaterally as a 2-mm spherical ROI surrounding peak glucose-responsive voxels (65). ASL data were motion corrected and then tagged and untagged images were subtracted to obtain perfusion-weighted images (64). ASL volumes were registered to the individual participant’s T_**1**_-weighted high-resolution anatomical volume using an affine registration with 12 degrees of freedom. Atlas ROIs where then inverse-transformed to individual native space and regional CBF was extracted for each ROI and participant.

### Statistical Analysis

Comparisons were conducted to measure FGF21 responses following both fructose vs glucose and sucrose vs glucose ingestion. The primary outcomes of interest were FGF21 responses after (1) fructose vs glucose (Experiment 1); (2) sucrose vs glucose consumption (Experiment 2); (3) associations between high vs low dietary added sugar intake and circulating FGF21 following sucrose ingestion; (4) associations between brain CBF and FGF21 responses to sucrose consumption; (5) effects of BMI status on the above outcomes. We used linear mixed-effects regression models, adjusting for age (66) and sex (20), with a random intercept for drink randomization order. For models examining associations between FGF21 and neural response to sucrose, we additionally adjusted for mean CBF (mCBF) and insulin AUC_35_, and glucose AUC_35_ to account for potential effects of circulating insulin and/or glucose on neural signaling. For longitudinal models that included repeated measurements over time, a random intercept for subject was included with an unstructured covariance matrix. In Experiment 1, we calculated FGF21 response to fructose and glucose as a change score of the value at ∼75min minus value at baseline. In Experiment 2, FGF21 response to sucrose and glucose were calculated as at the change score of the value at 120min vs. baseline. Additionally, in Experiment 2, we calculated total area under the curve (AUC)_35_ for plasma FGF21 and CBF in each ROI using the trapezoid method (also known as the trapezoidal rule (TR)) (67–70). TR AUC is calculated by adding the areas under the graph between each pair of consecutive observations, as previously described (67, 70). We used AUC across the first phase 35min testing period (AUC_35_) because this time frame was common to both the neuroimaging and blood sampling testing periods, and this allowed us to examine the neural and endocrine responses that were temporally related. Furthermore, in Experiment 2, dietary added sugar intake (i.e., low or high) between the BMI groups was compared using analysis of variance. We found FGF21 values to be right-skewed; for statistical analyses, both FGF21 raw data and AUC values were cubic root transformed to better meet the assumptions of normality. P-values <0.05 were interpreted as statistically significant. SAS 9.4 statistical software (SAS Institute, Cary, NC USA) was used for all data analyses, with regression models performed in PROC MIXED.

## Acknowledgements

The authors would like to thank the volunteers who participated in this study. The authors would also like to thank Ana Romero, Enrique Trigo, Reina Maniego, Hilary Dorton, Esther Jahng, Brandon Ge, Lloyd Nate Overholtzer, Jada Hislop, Martina Erdstein, and Priyanka Dave for assisting with study visits and recruiting volunteers, and the staff at the Dornsife Cognitive Neuroimaging Center and Diabetes and Obesity Research Institute of the University of Southern California, especially Lilit Baronikian for running the metabolic assays.

## Author Contributions

Conceptualization, K.A.P.; Methodology, K.A.P.; Supervision, K.A.P.; Funding Acquisition, K.A.P; Project Administration, A.G.Y., B.A., A.D.; Investigation, A.G.Y., B.A., A.D.; Formal Analysis, J.M.A. and T.A.P.; Writing – Original Draft, A.G.Y. and K.A.P.; Visualization, A.G.Y. and K.A.P.; Writing – Review & Editing, all authors.

## Data Availability

The datasets generated and analyzed during the current study are available from the corresponding author (K.A.P.) on reasonable request, and all brain imaging data are available in online digital repository (57).

