## Supplemental Information for "FGF21 Response Varies by Sugar Type and is Associated with Body Weight, Dietary Added Sugar, and Neural Signaling in Humans"

**This PDF file includes:**

Tables S1 to S5

Figure S1

**Table S1.** Least squares means for FGF21 response to fructose and glucose ingestion (Experiment 1).

| Whole Cohort | | | |
|---|---|---|---|
| **Drink** | **LS Mean (SE)** | **95% CI** | **p** |
| Fructose | 4.09 (0.28) | 3.54 to 4.65 | 0.01* |
| Glucose | 3.28 (0.27) | 2.73 to 3.82 | |
| **Healthy Weight** | | | |
| **Drink** | **LS Mean (SE)** | **95% CI** | **p** |
| Fructose | 4.05 (0.37) | 3.29 to 4.82 | 0.10 |
| Glucose | 3.23 (0.36) | 2.49 to 3.98 | |
| **Obesity** | | | |
| **Drink** | **LS Mean (SE)** | **95% CI** | **p** |
| Fructose | 4.14 (0.42) | 3.25 to 5.03 | 0.03* |
| Glucose | 3.35 (0.42) | 2.46 to 4.24 | |

FGF21 response calculated as values at ~75 minutes minus baseline. FGF21 values were cubic root transformed and LS means show FGF21 values adjusted for age and sex. *Indicates p values (which depict between drink difference) are statistically significant at $p<0.05$.

**Table S2.** Least squares means for FGF21 response to sucrose and glucose ingestion (Experiment 2).

| Whole Cohort | | | |
|---|---|---|---|
| **Drink** | **LS Mean (SE)** | **95% CI** | **p** |
| **Sucrose** | 6.26 (0.20) | 5.86 to 6.66 | <0.01* |
| **Glucose** | 2.54 (0.20) | 2.14 to 2.94 | |
| **Healthy Weight** | | | |
| **Drink** | **LS Mean (SE)** | **95% CI** | **p** |
| **Sucrose** | 5.87 (0.34) | 5.16 to 6.58 | <0.01* |
| **Glucose** | 2.36 (0.35) | 1.64 to 3.08 | |
| **Overweight** | | | |
| **Drink** | **LS Mean (SE)** | **95% CI** | **p** |
| **Sucrose** | 6.33 (0.37) | 5.57 to 7.09 | <0.01* |
| **Glucose** | 2.35 (0.36) | 1.60 to 3.10 | |
| **Obesity** | | | |
| **Drink** | **LS Mean (SE)** | **95% CI** | **p** |
| **Sucrose** | 6.61 (0.36) | 5.85 to 7.37 | <0.01* |
| **Glucose** | 2.92 (0.37) | 2.15 to 3.68 | |

FGF21 response calculated as values at 120 minutes minus baseline. FGF21 values were cubic root transformed and LS means show FGF21 values adjusted for age and sex. *Indicates p values (which depict between drink difference) are statistically significant at $p<0.05$.

**Table S3.** Least squares means for change in circulating FGF21 levels following sucrose ingestion stratified by BMI status and dietary added sugar intake (Experiment 2).

| BMI Status | Added Sugar Intake | LS Mean (SE) | p | N |
|---|---|---|---|---|
| Healthy Weight | Low | 5.33 (0.46) | 0.02* | 15 |
| | High | 7.23 (0.55) | | 10 |
| Overweight | Low | 6.02 (0.69) | 0.48 | 12 |
| | High | 6.75 (0.72) | | 11 |
| Obesity | Low | 6.09 (0.46) | 0.14 | 15 |
| | High | 7.59 (0.78) | | 6 |

*indicates p values are statistically significant at p<0.05. P values adjusted for age and sex. FGF21 response calculated as values at 120 minutes minus baseline. FGF21 values were cubic root transformed.

**Table S4.** Associations between FGF21 and cerebral blood flow (CBF) $AUC_{35}$ in a priori regions-of-interest (ROI) following sucrose ingestion among whole cohort (Experiment 2).

| Brain ROI | β Estimate | 95% CI | p |
|---|---|---|---|
| **Dorsal Striatum** | -7.66 | -14.48 to -0.85 | 0.03* |
| **Hippocampus** | -5.33 | -14.05 to 3.40 | 0.24 |
| **Hypothalamus** | 0.19 | -8.29 to 8.68 | 0.96 |
| **Insula** | -7.69 | -18.13 to 2.76 | 0.15 |
| **Nucleus Accumbens** | -10.29 | -21.61 to 1.03 | 0.08 |

*indicates p values are statistically significant at $p<0.05$. P values adjusted for age, sex, BMI status, mean cerebral blood flow (mCBF), insulin $AUC_{35}$, and glucose $AUC_{35}$. FGF21 values were cubic root transformed.

**Table S5.** Associations between FGF21 and cerebral blood flow (CBF) $AUC_{35}$ in a priori regions-of-interest (ROI) following sucrose ingestion, stratified by BMI status (Experiment 2).

| Brain ROI | BMI Status | β Estimate | 95% CI | p |
|---|---|---|---|---|
| **Dorsal Striatum** | Healthy Weight | -15.70 | -25.98 to -5.41 | 0.01* |
| | Overweight | -4.00 | -19.71 to 11.72 | 0.62 |
| | Obesity | -12.45 | -27.42 to 2.51 | 0.13 |
| **Hippocampus** | Healthy Weight | -20.60 | -36.79 to -4.42 | 0.02* |
| | Overweight | -17.87 | -36.93 to 1.19 | 0.08 |
| | Obesity | 1.34 | -14.37 to 17.05 | 0.87 |
| **Hypothalamus** | Healthy Weight | -4.30 | -19.19 to 10.59 | 0.58 |
| | Overweight | -11.44 | -34.08 to 11.20 | 0.34 |
| | Obesity | -8.87 | -24.68 to 6.94 | 0.29 |
| **Insula** | Healthy Weight | -16.54 | -32.62 to -0.47 | 0.06 |
| | Overweight | -7.35 | -35.00 to 20.30 | 0.60 |
| | Obesity | -7.43 | -29.51 to 14.66 | 0.52 |
| **Nucleus Accumbens** | Healthy Weight | -13.25 | -33.88 to 7.39 | 0.23 |
| | Overweight | -19.56 | -43.43 to 4.31 | 0.13 |
| | Obesity | -11.04 | -42.11 to 20.04 | 0.50 |

*indicates p values are statistically significant at $p<0.05$. P values adjusted for age, sex, mean cerebral blood flow (mCBF), insulin $AUC_{35}$, and glucose $AUC_{35}$. FGF21 values were cubic root transformed.

**Figure S1.** Trajectories for post-ingestive plasma FGF21 responses.

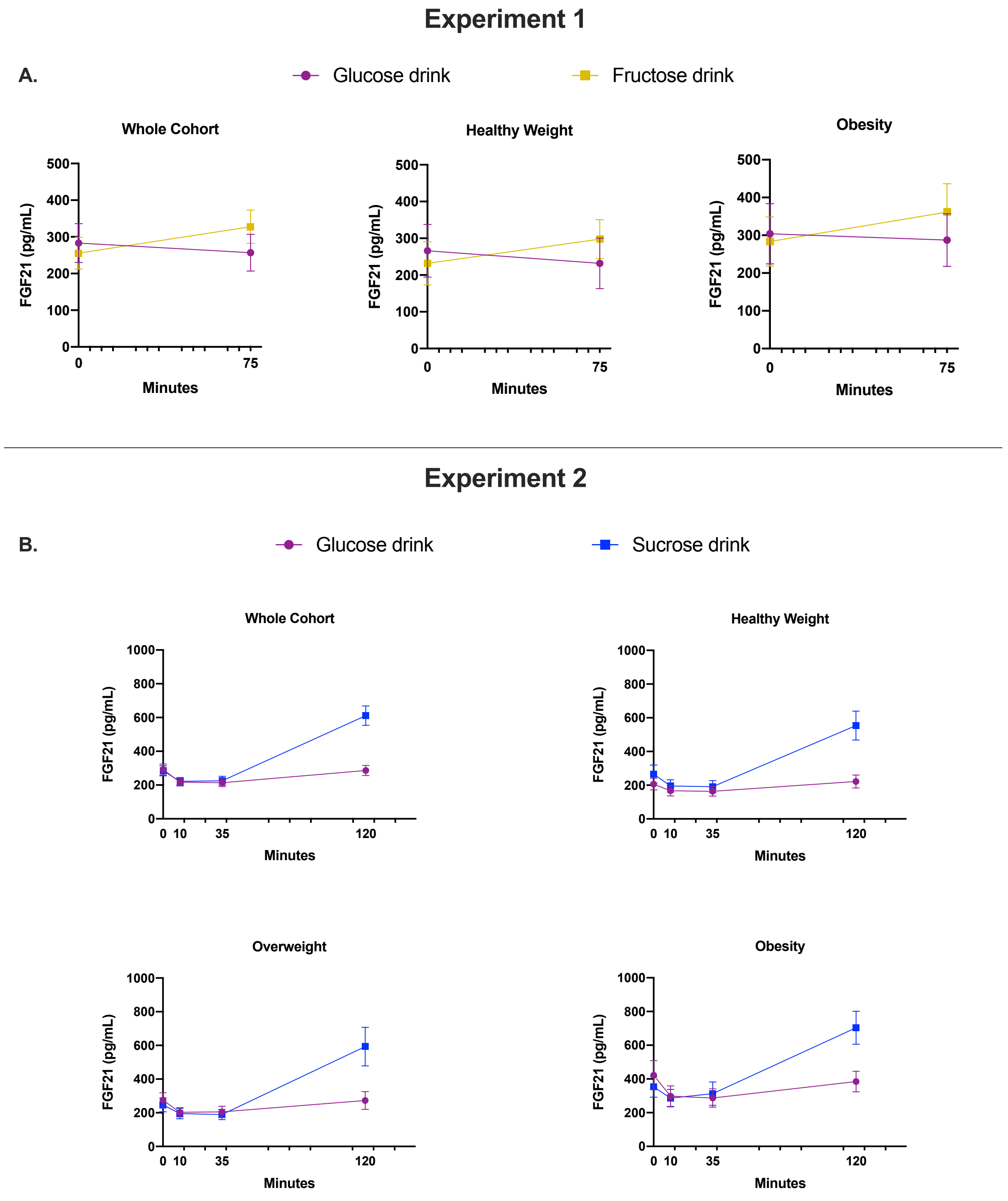

Following acute A) fructose and glucose (Experiment 1, N=38) at 0 and 75 minutes; and B) sucrose and glucose consumption (Experiment 2, N=69) at 0, 10, 35, and 120 minutes, among the cohort as a whole and stratified by body mass index (BMI) groups. Data are expressed as raw/unadjusted mean ± SEM for visual purposes, but all statistical analyses were based on cubic root transformed FGF21 values and adjusted for covariates. See also Figure 1 and Tables S1 and S2.
